# Dendritic Wave Recurrent Neural Networks

**DOI:** 10.64898/2026.07.03.736415

**Authors:** Yoshimasa Kubo

**Affiliations:** Department of Computer Science, Lakehead University, Thunder Bay, Canada

## Abstract

Wave recurrent neural networks (wRNNs) are biologically inspired recurrent architectures that use traveling-wave dynamics to support sequence learning and memory. However, their input-to-hidden pathway remains relatively simple compared with biological neurons, where dendrites perform nonlinear input integration. In this study, we introduce the Dendritic Wave Recurrent Neural Network (DW-RNN), which augments the input pathway of the wRNN with nonlinear basal dendritic branches while preserving the original recurrent wave dynamics. We evaluate DW-RNN on a simple copy task, sequential MNIST (sMNIST), permuted sequential MNIST (psMNIST), and noisy sequential CIFAR-10 (nsCIFAR-10). On the copy task, DW-RNN shows learning behavior comparable to the standard wRNN, suggesting that dendritic input integration does not disrupt the recurrent wave-based memory mechanism. On the three sequential image-classification benchmarks, DW-RNN outperforms the standard wRNN, improving accuracy from 97.27 ± 0.15% to 97.82 ± 0.12% on sMNIST, from 96.74 ± 0.17% to 96.92 ± 0.10% on psMNIST, and from 54.30 ± 0.79% to 55.65 ± 0.55% on nsCIFAR-10. In addition to improving mean accuracy, DW-RNN exhibits lower across-seed variability on all three classification benchmarks, suggesting that dendritic input integration may improve the stability of wRNN training. Hidden-activity visualizations further show that DW-RNN preserves the characteristic traveling-wave patterns of the original wRNN. These results suggest that dendritic computation and traveling-wave recurrent dynamics provide complementary mechanisms for biologically inspired sequence learning.

## 1 Introduction

Wave recurrent neural networks (wRNNs) are biologically inspired recurrent neural networks motivated by neural oscillations and the spatial organization of neural activity in the brain [Keller et al., 2024]. Previous work showed that wRNNs achieve strong performance on sequential image-classification tasks and memory-based tasks such as the copy task, outperforming or competing with conventional recurrent architectures such as long short-term memory networks (LSTMs) [Hochreiter and Schmidhuber, 1997] and gated recurrent units (GRUs) [Chung et al., 2014]. Importantly, the hidden-layer dynamics of wRNNs exhibit traveling-wave activity, resembling wave-like neural activity observed in the brain. Since traveling waves have been implicated in temporal processing and memory, these results suggest that wave-like recurrent dynamics may provide a useful computational mechanism for sequence learning.

Another class of biologically inspired neural-network models is dendritic neural networks [Grewal et al., 2021, Chavlis and Poirazi, 2025, Guerguiev et al., 2017, Kubo, 2026]. Dendrites play an important role in nonlinear input integration and can amplify or attenuate neuronal activity. Prior studies have shown that incorporating dendritic mechanisms, particularly basal dendritic mechanisms for input integration, into artificial neural networks can improve performance in settings such as continual learning [Grewal et al., 2021] and reduce overfitting [Chavlis and Poirazi, 2025]. Recent work has also shown that dendritic recurrent neural networks trained with biologically plausible learning algorithms, such as equilibrium propagation [Scellier and Bengio, 2017, 2019, Ernoult et al., 2019, Laborieux et al., 2021, Laborieux and Zenke, 2022] combined with predictive learning rules [Luczak et al., 2022, Luczak and Kubo, 2022, Kubo et al., 2023], can achieve performance competitive with backpropagation-trained dendritic networks [Kubo, 2026]. In that study, both basal dendritic mechanisms for feedforward input integration and apical dendritic mechanisms for feedback integration were incorporated into recurrent neural networks. These findings suggest that dendritic computation may provide a useful mechanism for improving recurrent neural networks while maintaining biological plausibility.

In this study, we introduce the Dendritic Wave Recurrent Neural Network (DW-RNN), which augments the input-to-hidden pathway of the wRNN with nonlinear basal dendritic branches. The proposed model is inspired by recent dendritic neural-network models [Kubo, 2026], but preserves the recurrent wave dynamics of the original wRNN. This design allows us to examine whether nonlinear dendritic input integration can improve sequence learning without disrupting the traveling-wave recurrent mechanism that supports temporal processing in wRNNs.

We evaluate DW-RNN on a simple copy task [Graves et al., 2014], sequential MNIST (sMNIST) [Lamb et al., 2016], permuted sequential MNIST (psMNIST) [Arjovsky et al., 2016], and noisy sequential CIFAR-10 (nsCIFAR-10) [Chang et al., 2019]. On the copy task, DW-RNN shows learning behavior comparable to the standard wRNN, suggesting that basal dendritic input integration does not disrupt the recurrent wave-based memory mechanism. On the three sequential image-classification benchmarks, DW-RNN outperforms the standard wRNN, suggesting that nonlinear dendritic input integration improves sequence learning. In addition to improving mean accuracy, DW-RNN shows lower standard deviation across random seeds on all three classification benchmarks, suggesting reduced sensitivity to initialization and improved training stability. We further analyze hidden-layer activations to determine whether the proposed dendritic extension preserves the traveling-wave dynamics of the original wRNN. Our results show that DW-RNN improves performance while maintaining characteristic wave-like hidden activity, indicating that dendritic input integration can enhance recurrent sequence models without disrupting their underlying wave dynamics.

## 2 Methods

### 2.1 Wave Recurrent Neural Networks (wRNNs)

Wave Recurrent Neural Networks (wRNNs) were introduced as a simple recurrent neural-network architecture designed to generate traveling-wave dynamics in the hidden layer [Keller et al., 2024]. The model is motivated by observations that traveling waves of neural activity are widely observed in the brain and have been associated with temporal processing and memory. In the wRNN, wave-like hidden dynamics provide an inductive bias for encoding recent sequential inputs, allowing the hidden state to act as a wave-field memory buffer.

A standard simple recurrent neural network updates its hidden state using a fully connected recurrent transformation. In contrast, the wRNN replaces this dense recurrent transformation with a circular one-dimensional convolution over the hidden state. The hidden-state update is given by

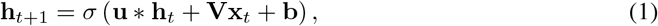

where **h**_*t*_ is the hidden state at time step *t*, **x**_*t*_ is the input, **u** * **h**_*t*_ denotes circular convolution over the hidden dimension, **V** is the input-to-hidden weight matrix, **b** is a bias term, and *σ*(·) is the hidden-state activation function. The recurrent convolution can be interpreted as local connectivity on a ring of hidden units, allowing activity to propagate across hidden positions over time.

Previous work showed that this architecture produces clear diagonal bands of activity in the hidden layer, corresponding to traveling waves propagating through the hidden state [Keller et al., 2024]. These wave dynamics were shown to improve learning on synthetic memory tasks and sequential image-classification benchmarks, including sequential MNIST, permuted sequential MNIST, and noisy sequential CIFAR-10. In particular, wRNNs were reported to learn faster than comparable wave-free recurrent networks and to achieve competitive performance with more complex recurrent architectures such as LSTMs and GRUs.

In the present study, we build directly on the wRNN architecture. Our goal is not to modify the recurrent wave mechanism itself, but rather to improve how external inputs are integrated into the wave-based hidden state. Therefore, in the proposed Dendritic Wave Recurrent Neural Network (DW-RNN), the recurrent convolutional term **u** * **h**_*t*_ is preserved, while the standard linear input term **Vx**_*t*_ is replaced with nonlinear basal dendritic input integration. This modification is described in the next subsection.

### 2.2 Dendritic Neurons

In this study, we extend the Wave Recurrent Neural Network (wRNN) by incorporating nonlinear basal dendritic branches into the input-to-hidden pathway. The dendritic-neuron formulation is based primarily on our previous dendritic recurrent neural-network model [Kubo, 2026], in which each neuron is represented as a collection of nonlinear dendritic branches whose outputs are integrated before the somatic activation. More broadly, this design is motivated by prior work showing that dendritic compartments can provide structured nonlinear processing in artificial neural networks [Grewal et al., 2021, Chavlis and Poirazi, 2025, Guerguiev et al., 2017].

In biological neurons, basal dendrites primarily receive feedforward inputs, whereas apical dendrites are often associated with feedback and contextual signals. The previous dendritic recurrent model incorporated both basal dendritic mechanisms for feedforward input integration and apical dendritic mechanisms for feedback integration [Kubo, 2026]. In contrast, the present study focuses specifically on modifying the feedforward input pathway of wRNNs while preserving their recurrent wave dynamics. Therefore, DW-RNN uses only basal dendritic mechanisms to replace the standard linear input term **Vx**_*t*_.

In the original wRNN, the hidden state is updated by combining a linear input term, **Vx**_*t*_, with a recurrent circular convolution term, **u** * **h**_*t*_, which produces traveling-wave dynamics. In the proposed Dendritic Wave Recurrent Neural Network (DW-RNN), we replace the standard linear input term **Vx**_*t*_ with a nonlinear dendritic input term **d**(**x**_*t*_), while leaving the recurrent wave convolution unchanged. This design allows us to examine whether dendritic input integration can improve sequence learning without disrupting the recurrent mechanism responsible for wave propagation.

For each hidden unit, the input signal is processed by multiple basal dendritic branches. Each branch receives a subset of input features, applies a linear transformation followed by a nonlinear activation, and produces a local dendritic response. Let *z*_*i,k,t*_ denote the response of the *k*-th basal branch associated with hidden unit *i* at time step *t*. The branch response is computed as

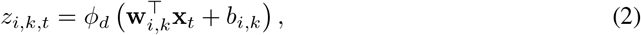

where **x**_*t*_ is the input at time step *t*, **w**_*i,k*_ and *b*_*i,k*_ are the branch weight vector and bias, and *ϕ*_*d*_(·) denotes the dendritic branch nonlinearity.

The dendritic input to hidden unit *i* is obtained by averaging the outputs of its *K* basal branches:

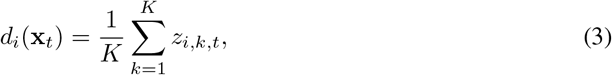

where *K* is the number of basal dendritic branches per hidden unit. In vector form, the dendritic input to the hidden layer is denoted by **d**(**x**_*t*_).

The DW-RNN hidden-state update is then defined as

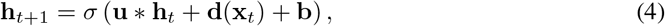

where **h**_*t*_ is the hidden state, **u** * **h**_*t*_ denotes the recurrent circular convolution used in the original wRNN, **b** is a bias term, and *σ*(·) is the hidden-state activation function. Importantly, the recurrent convolutional pathway is unchanged from the original wRNN. Thus, DW-RNN modifies only the input-to-hidden pathway, replacing the linear term **Vx**_*t*_ with nonlinear basal dendritic input integration, while preserving the traveling-wave recurrent dynamics.

We used different dendritic nonlinearities depending on the task. For the copy task, sMNIST, and psMNIST, we used a gain-controlled hyperbolic tangent nonlinearity,

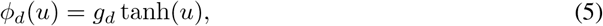

where *g*_*d*_ controls the strength of the dendritic branch response. For the main sMNIST and psMNIST experiments, we used *g*_*d*_ = 4. For noisy CIFAR-10, we instead used a leaky-ReLU dendritic nonlinearity,

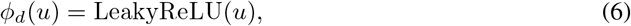

without an additional dendritic gain parameter. This choice provided more stable training for the higher-dimensional noisy visual sequence.

In our experiments, the number of basal branches, branch sparsity, and dendritic nonlinearity were treated as hyperparameters. Overall, this formulation introduces structured sparse nonlinear input processing into wRNNs while maintaining the original recurrent wave mechanism.

### 2.3 Model and Dataset Specification

We first evaluated DW-RNN and the standard wRNN on a copy task, which has been widely used to assess sequence memory in recurrent neural networks [Keller et al., 2024, Graves et al., 2014, Arjovsky et al., 2016, Gu et al., 2020]. In this task, the model receives an initial sequence of 10 symbols, followed by a delay period of length *T*_delay_, a delimiter symbol, and additional blank symbols. The target output is zero at all time steps except for the final 10 steps, where the model must reproduce the initial symbol sequence. Longer delay lengths require the model to retain information for a longer period and therefore make the task more difficult. We evaluated both models using *T*_delay_ ∈ {0, 30, 80}.

We then evaluated DW-RNN and wRNN on three sequential image-classification benchmarks: sequential MNIST (sMNIST), permuted sequential MNIST (psMNIST), and noisy sequential CIFAR-10 (nsCIFAR-10). These tasks test the ability of recurrent models to process long input sequences while maintaining task-relevant information over time.

The hyperparameters used for each task are summarized in Table 1. Unless otherwise specified, the same hyperparameters were used for DW-RNN and wRNN to ensure a fair comparison.

**Table 1:** Hyperparameters used for the copy task and sequential image-classification tasks. The same settings were used for wRNN and DW-RNN unless otherwise noted. Here, *α* denotes the learning rate.

| Hyperparameter | Copy | sMNIST | psMNIST | nsCIFAR-10 |
| --- | --- | --- | --- | --- |
| Learning rate $\alpha$ | $1 \times 10^{-3}$ | $1 \times 10^{-4}$ | $1 \times 10^{-4}$ | $1 \times 10^{-4}$ |
| Hidden size | 100 | 256 | 256 | 256 |
| Batch size | 128 | 128 | 128 | 128 |
| Channels | 6 | 16 | 16 | 16 |
| Hidden activation | ReLU | ReLU | ReLU | ReLU |
| Filter size | 3 | 3 | 3 | 3 |

For the dendritic input pathway, we used a fixed configuration across all datasets, consisting of two basal branches per hidden unit and a branch sparsity of 0.5. These settings were chosen to balance model expressivity and computational efficiency. For the nonlinear transformation within dendritic branches, we used a hyperbolic tangent nonlinearity for the copy task, sMNIST, and psMNIST. For nsCIFAR-10, we used a leaky-ReLU dendritic nonlinearity, which provided more stable training for the higher-dimensional noisy visual sequence.

All models were trained using the Adam optimizer [Kingma and Ba, 2014] with cross-entropy loss. Each experiment was repeated across five random seeds, and we report the mean and standard deviation of the corresponding performance metrics. Following the training protocol of the original wRNN study [Keller et al., 2024], we used gradient clipping with a maximum norm of 1.0. For the sequential image-classification tasks, the learning rate was reduced by a factor of 10 every 100 epochs.

#### Code availability

The code used in this study is currently being organized and will be made available in a public GitHub repository. The repository link will be added in a revised version of this preprint.

## 3 Results

### 3.1 Copy Task

Figure 1 shows the learning curves of DW-RNN and the standard wRNN on the copy task for delay lengths *T*_delay_ ∈ {0, 30, 80}. Both models quickly learned the task across all tested delay lengths, including the longest delay condition of *T*_delay_ = 80. DW-RNN exhibited learning behavior comparable to the standard wRNN, indicating that replacing the linear input-to-hidden pathway with nonlinear basal dendritic input integration does not impair the sequence-memory capability of wRNNs. These results suggest that the proposed dendritic extension preserves the memory performance of the original wRNN, while allowing DW-RNN to benefit from nonlinear input integration on the sequential image-classification tasks.

**Figure 1.**
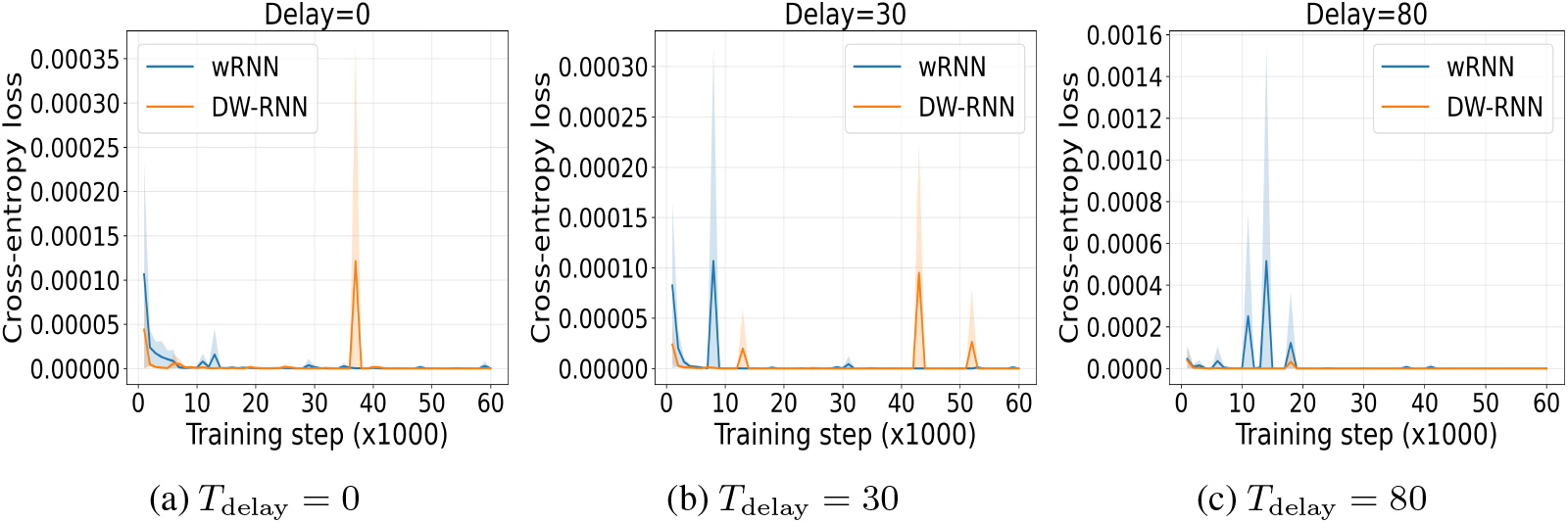
Learning curves on the copy task for delay lengths *T*_delay_ ∈ {0, 30, 80}. The memory length was fixed to 10 symbols. Both wRNN and DW-RNN were trained using cross-entropy loss for categorical token prediction. Solid lines indicate the mean over five random seeds, and shaded regions indicate standard deviation.

### 3.2 sMNIST

The first column of Table 2 reports the performance of wRNN and DW-RNN on sMNIST. DW-RNN achieved an accuracy of 97.82 ± 0.12%, compared with 97.27 ± 0.15% for the standard wRNN, corresponding to an improvement of 0.55 percentage points.

**Table 2:** Test accuracy (%) on sequential image-classification tasks. Results are reported as mean ± standard deviation over five random seeds. DW-RNN improves mean accuracy and reduces across-seed variability on all three benchmarks.

| Model | sMNIST | psMNIST | nsCIFAR-10 |
| --- | --- | --- | --- |
| wRNN | $97.27 \pm 0.15$ | $96.74 \pm 0.17$ | $54.30 \pm 0.79$ |
| DW-RNN | **$97.82 \pm 0.12$** | **$96.92 \pm 0.10$** | **$55.65 \pm 0.55$** |
| Improvement | +0.55 | +0.18 | +1.35 |

Figure 2a shows the corresponding learning curves. DW-RNN converged faster than the standard wRNN and reached a higher final accuracy, suggesting that nonlinear basal dendritic input integration improves learning on the standard sequential MNIST task.

**Figure 2.**
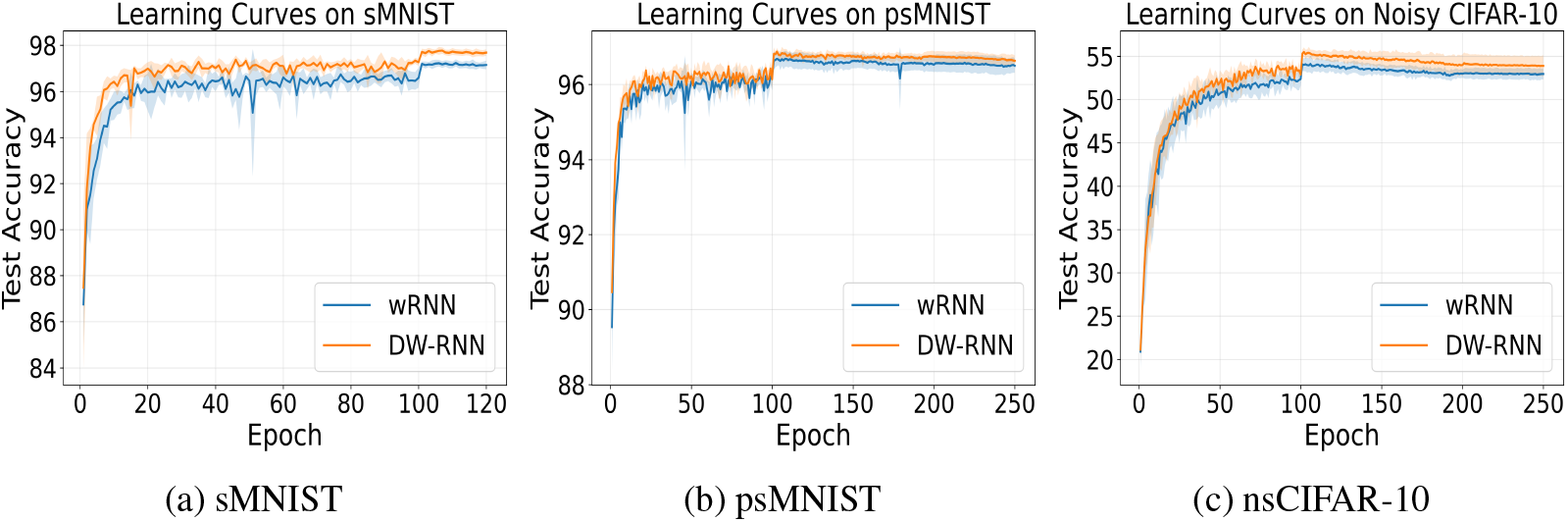
Learning curves on sequential image-classification tasks.

### 3.3 psMNIST

The second column of Table 2 summarizes the results on psMNIST. DW-RNN achieved an accuracy of 96.92 ± 0.10%, while the standard wRNN achieved 96.74 ± 0.17%. Although the improvement is smaller than that observed on sMNIST, DW-RNN still improved performance by 0.18 percentage points and showed lower across-seed variability.

As shown in Figure 2b, DW-RNN also converged faster than the standard wRNN on psMNIST. This suggests that dendritic input integration remains beneficial even when the temporal structure of the input sequence is disrupted by a fixed random permutation.

### 3.4 nsCIFAR-10

The final column of Table 2 presents the results on nsCIFAR-10. DW-RNN achieved an accuracy of 55.65 ± 0.55%, compared with 54.30 ± 0.79% for the standard wRNN. This corresponds to an improvement of 1.35 percentage points, which is the largest improvement among the three sequential image-classification benchmarks.

Figure 2c shows that DW-RNN again converged faster than the standard wRNN. These results suggest that nonlinear basal dendritic input integration is particularly useful for the higher-dimensional noisy visual sequence used in nsCIFAR-10.

### 3.5 Wave Comparisons

The copy-task results show that DW-RNN preserves the sequence-memory capability of the standard wRNN, while the classification results show that DW-RNN improves performance on sequential image-classification tasks. However, these results alone do not indicate whether the proposed dendritic extension preserves the characteristic traveling-wave dynamics of the original wRNN. To examine this, we visualized the hidden-layer activations of DW-RNN and wRNN on sMNIST.

Figure 3 shows representative hidden-layer activity patterns for both models. In both DW-RNN and wRNN, the hidden activations exhibit clear diagonal wave-like structures over time, indicating that activity propagates across hidden positions during sequence processing. These results suggest that nonlinear basal dendritic input integration does not disrupt the recurrent traveling-wave dynamics of wRNNs. Together with the improved classification performance, this indicates that dendritic input integration can enhance sequence learning while preserving the wave-like hidden dynamics of the original architecture.

**Figure 3.**
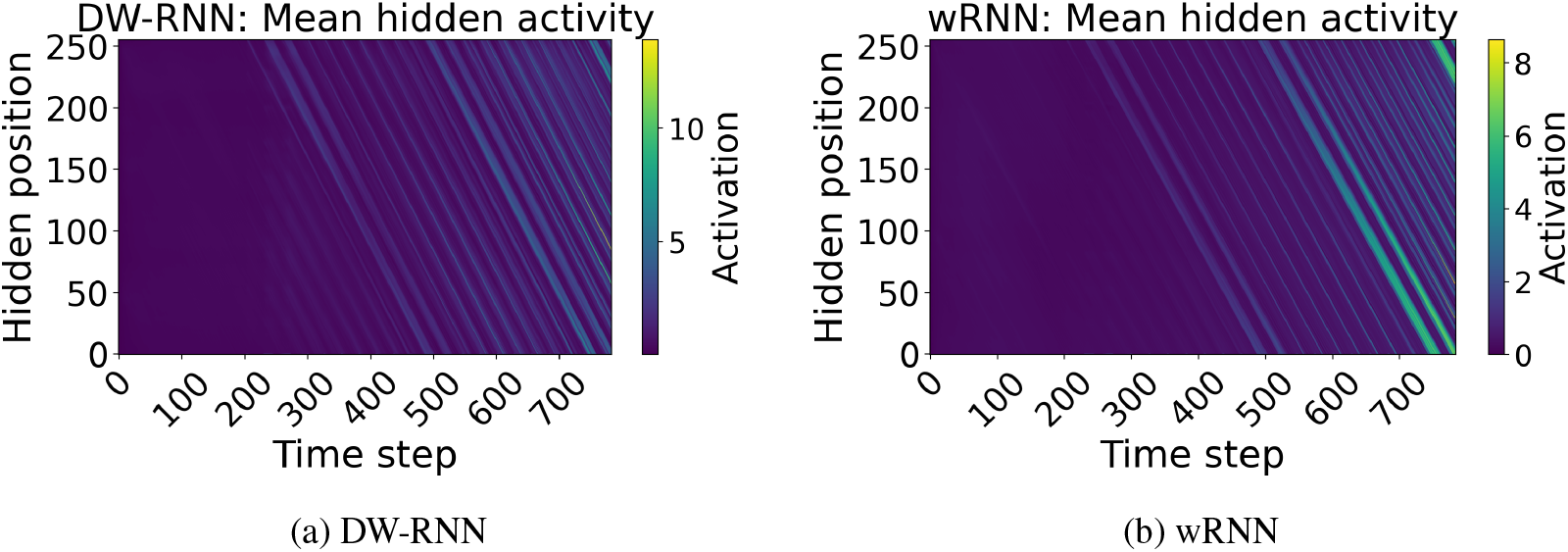
Hidden-layer activity on sMNIST for DW-RNN and the standard wRNN. Each heatmap shows the mean hidden activation across channels for a representative test example. The horizontal axis represents time step, and the vertical axis represents hidden position. DW-RNN preserves the characteristic diagonal wave-like activity patterns observed in the standard wRNN, indicating that nonlinear basal dendritic input integration does not disrupt the recurrent traveling-wave dynamics.

## 4 Discussion

In this study, we introduced the Dendritic Wave Recurrent Neural Network (DW-RNN), which incorporates nonlinear basal dendritic input integration into the Wave Recurrent Neural Network (wRNN). The dendritic mechanism was based on our previous dendritic neural-network model [Kubo, 2026], but was applied specifically to the input-to-hidden pathway of wRNNs while preserving the recurrent wave dynamics of the original architecture. Our results show that DW-RNN maintains the sequence-memory capability of the standard wRNN on the copy task, while improving performance on all three sequential image-classification benchmarks.

The main comparison in this study was between DW-RNN and the original wRNN, because our goal was to isolate the effect of adding nonlinear dendritic input integration to the wRNN architecture. Nevertheless, it is useful to compare our results with previously reported recurrent-network baselines. The original wRNN study [Keller et al., 2024] compared wRNNs with other recurrent architectures, including LSTMs [Hochreiter and Schmidhuber, 1997] and GRUs [Chung et al., 2014], using results reported in prior work [Chang et al., 2019]. On sMNIST, LSTM and GRU achieved accuracies of 98.8% and 99.1%, respectively, which remain higher than the performance of DW-RNN. However, on psMNIST, the reported LSTM and GRU accuracies were 92.9% and 94.1%, whereas DW-RNN achieved 96.92 ± 0.10%. Similarly, on nsCIFAR-10, the reported LSTM and GRU accuracies were 11.6% and 43.8%, whereas DW-RNN achieved 55.65 ± 0.55%. These comparisons suggest that DW-RNN is particularly effective on tasks requiring robust sequence processing under permutation or noise, while preserving the biologically inspired wave dynamics of the original wRNN.

An important implication of these results is that dendritic input integration and traveling-wave recurrent dynamics appear to provide complementary computational mechanisms. The recurrent wave dynamics of wRNNs support temporal processing and sequence memory, while nonlinear basal dendritic branches enrich the input-to-hidden transformation. The copy-task results suggest that adding dendritic branches does not impair the sequence-memory capability of the original wRNN. In contrast, the classification results show that dendritic input integration can improve performance and reduce across-seed variability on sequential image-classification tasks. Hidden-activity visualizations further show that DW-RNN preserves the characteristic wave-like activity patterns of wRNNs. Together, these findings suggest that dendritic computation can enhance wave-based recurrent models without disrupting their underlying hidden-state dynamics.

A limitation of the present study is that DW-RNN was trained using backpropagation through time (BPTT). Although the architecture is biologically inspired, BPTT is not generally considered biologically plausible. In future work, we aim to replace BPTT with equilibrium propagation (EP), a biologically plausible learning algorithm based on differences between free and nudged network states [Scellier and Bengio, 2017, 2019, Ernoult et al., 2019, Laborieux et al., 2021, Laborieux and Zenke, 2022]. We also plan to combine EP with predictive learning rules [Luczak et al., 2022, Luczak and Kubo, 2022, Kubo et al., 2023], which may further improve the biological plausibility of training in dendritic recurrent architectures.

Another promising future direction is to introduce heterogeneous temporal dynamics [Kubo et al., 2026, Perez-Nieves et al., 2021] into DW-RNN. In the present study, hidden units were updated using a common temporal discretization. However, biological neurons exhibit diverse temporal properties, including heterogeneous membrane time constants. Introducing heterogeneous update steps or time constants into DW-RNN may change the propagation speed, structure, and stability of traveling waves in the hidden layer. Such temporal heterogeneity could allow different hidden positions or channels to represent information over multiple time scales, which may be useful for long-range sequence modeling. Future work should examine how heterogeneous temporal dynamics interact with nonlinear dendritic input integration and whether they further improve memory, stability, and wave propagation.

Finally, DW-RNN may be useful beyond sequential image-classification tasks. Since wRNNs are designed to process long sequences through wave-like recurrent dynamics, and DW-RNN further improves nonlinear input integration, this architecture may be applicable to natural language processing and other sequence-modeling problems. Future work could evaluate DW-RNN on language-modeling or sequence-prediction tasks and compare it with standard recurrent baselines such as LSTMs and GRUs.

## Acknowledgments

This research was enabled in part by computational resources provided by the Digital Research Alliance of Canada (alliancecan.ca).

